# The Ataxin-2 protein is required in Kenyon cells for RNP-granule assembly and appetitive long-term memory formation

**DOI:** 10.1101/2025.05.15.654326

**Authors:** Camilla Roselli, Jens Hillebrand, Jenifer Kaldun, Vernon Leander Monteiro, Thomas Hurd, Simon G. Sprecher, Tamara Boto, Mani Ramaswami

**Affiliations:** Department of Genetics and Microbiology, Trinity College Dublin, Dublin, Ireland; Trinity College Institute of Neuroscience, Dublin, Ireland; Department of Physiology, School of Medicine, Trinity College Dublin, Dublin, Ireland; University of Fribourg, Department of Biology, Fribourg, Switzerland; National Center of Biological Sciences, Tata Institute of Fundamental Research, Bangalore, India; Department of Molecular Genetics, University of Toronto, Ontario, Canada; School of Physiology, Pharmacology and Neuroscience, University of Bristol, Bristol, United Kingdom

**Keywords:** long-term memory, appetitive, RNA-binding protein, translational regulation, neuronal granules, mRNP condensate

## Abstract

Ribonucleo-protein granules (mRNP granules) are thought to contribute to the control of neuronal mRNA translation required for consolidation of long-term memories. Consistent with this, the function of Ataxin-2 in mRNA granule assembly has been shown to be required for long-term olfactory habituation (LTH) in *Drosophila*, a form of non-associative memory. Knockdown of Ataxin-2 in either local interneurons (LNs) or projection neurons (PNs) of the insect antennal lobe disrupts LTH, leading to a model in which Ataxin-dependent translational control is required in both presynaptic and postsynaptic elements of the LN-PN synapse, whose potentiation has been causally linked to LTH. Here we use novel and established methods for cell-type specific perturbation to ask: (a) whether Ataxin-2 controls mRNA granule assembly in cell types beyond the few that have been examined; and (b) whether it functions not only in LTH, but also for long-term olfactory associative memory (LTM). We show that Ataxin-2 controls mRNP granule assembly in additional neuronal types, namely Kenyon Cells that encode associative memory, as well as more broadly in non-neuronal cells, e.g. in nurse cells in the egg chamber. Furthermore, selective knockdown of Atx2 in α/β and α’/β’ KCs blocks appetitive long-term but not short-term associative memories. Taken together these observations support a hypothesis that Ataxin-2 dependent translational control is widely required across different mnemonic circuits for consolidation of respective forms of long-term memories.

## INTRODUCTION

Long-term memories (LTMs) are distinguished from short-term memories (STMs) not only by their persistence across time, but also by the requirement for new protein synthesis in long-term synaptic plasticity that underlies LTM (Bailey et al., 1996). Several lines of evidence have suggested that activity-regulated synaptic translation of stored mRNAs contributes to long-term forms of plasticity and memory (Das et al., 2023; Donnelly et al., 2010; Martin & Kosik, 2002; Richter & Klann, 2009). Even though the requirement of *de novo* protein synthesis has been well established, specially using protein synthesis inhibitors (Agranoff et al., 1967; Hernandez & Abel, 2008; Squire & Barondes, 1972; Tully et al., 1994), some key questions remain open; for example: what are the translational mechanisms involved in such process? How different are these across cell types? And what is the diversity of cell types in which activity-dependent mRNA translation, and hence a specific type of plasticity, must occur for any particular form of long-term memory.

One of the mechanisms by which *de novo* protein synthesis can be regulated is via activity-dependent modulation of ribonucleoprotein granules (mRNP granules), dynamic, membrane-less organelles, which sequester translationally repressed mRNAs together with proteins that control their translation, localization and turnover (Buchan, 2014).

Work in the model organism *Drosophila melanogaster* has shown that several components of the neuronal-granule associated, translational-control machinery, such as: Pumilio (Vessey et al., 2006), Staufen (Barbee et al., 2006), EIF5C (Dubnau et al., 2003), Gld2 (Kwak et al., 2008), Orb2 (Hervás et al., 2016; Keleman et al., 2007; Majumdar et al., 2012; Mastushita-Sakai et al., 2010), FMRP and components of the miRNA pathway (Bolduc et al., 2008; Sudhakaran et al., 2013) are essential for consolidation of LTM under conditions where they are entirely dispensable for STM. Significant among these RNA-binding protein components of neuronal granules, is Ataxin-2 (Atx2), a protein altered in neurodegenerative disease, that can serve as both an activator or repressor of translation (Ciosk et al., 2004; Inagaki et al., 2020; Lee et al., 2018; Lim & Allada, 2013; Zhang et al., 2013, 2013) Atx2 is also unique in being required for the presence of a specific class of Me31B/DDX6 positive neuronal mRNP condensates, at least in inhibitory local interneurons (LNs) and excitatory projection neurons (PNs) of the *Drosophila* antennal lobes, where Atx2 must be present for olfactory LTH (Bakthavachalu et al., 2018; McCann et al., 2011; Sudhakaran et al., 2013).

The formation of mRNP granules is facilitated by multivalent interactions between mRNA molecules, the binding of RNA binding proteins (RBPs) to their target mRNAs as well as RBP interactions among each other particularly through intrinsically disordered regions (Protter & Parker, 2016). A detailed structure function analysis of the *Drosophila* Atx2 protein domains in cultured cells and *in vivo* identified a C-terminally located, intrinsically disordered region (cIDR) of the protein as being required for efficient neuronal granule formation, but dispensable for animal survival (Bakthavachalu et al., 2018). Significantly, similar to flies with reduced levels of Atx2, flies genetically engineered to lack only the Atx2 cIDR showed defective LTH but normal STH (Bakthavachalu et al., 2018; McCann et al., 2011; Sudhakaran et al., 2013). These data argue for a causal role for mRNP granules in the regulation of translational processes that underly long-term habituation (Bakthavachalu et al., 2018).

Neuronal mRNP condensates are not only of interest for their function in long-term memory, but also for their involvement in neurodegenerative diseases – particularly Amyotrophic Lateral Sclerosis (ALS) and frontotemporal dementia (FTD) - which typically show cell-type specificity (Ling et al., 2013; Nedelsky & Taylor, 2022). Polyglutamine expansions in human Atx2 protein, which increase the stability of Atx2 condensates (Wijegunawardana et al., 2024), are associated with spinocerebellar ataxia-2 (SCA2) and Amyotrophic Lateral Sclerosis (ALS) (Elden et al., 2010). Remarkably cytotoxicity seen in fly models of ALS, is greatly reduced in animals lacking the Atx2 cIDR (Bakthavachalu et al., 2018) directly connecting mRNP granules function with disease progression. Further, the involvement of Atx2 in LTM has as well been observed in human patients. For instance, SCA2 patients, with a mutation in the human Atx2 locus, also show a decline in verbal memory (Rodríguez-Labrada et al., 2019).

To more deeply examine Atx2’s functions *in vivo*, we investigated (a) whether Atx2 contributes to the assembly of mRNP condensates in multiple neuronal and non-neuronal cells; and (b) how and how widely mRNP granules contribute to associative long-term memory, which extensive studies have shown to depend on plasticity of a different subset of neurons: namely, in synapses made by Kenyon cells (KCs) onto output neurons (MBONs) of the *Drosophila* mushroom body, a structure known to be the major site of associative learning and memory in insects (Campbell & Turner, 2010).

## MATERIAL AND METHODS

### Fly lines and husbandry

All genotypes were raised on standard cornmeal medium under LD (12 hr Light: 12 hr Dark) cycles at 25°C. For behavioural analysis the flies were grown at 18°C until eclosure, then the animals were moved to 30°C for a week for RNAi expression. The following Gal4 and effector lines were used: *MB247-gal4*, *c739-gal4*, *c305a-gal4*, *5-HTR1b-gal4, UAS-dcr* (S. Sprecher lab stock), *nos-gal4::VP16* (RRID:BDSC 4937), *tubGal80(ts)* (RRID:BDSC 7018), *UAS-Atx2 RNAi*^1^ (VDRC 34955), *UAS-Atx2 RNAi^2^*(RRID:BDSC 36114), *UAS-Atx2 RNAi^3^* (RRID:BDSC 67878), *hsFLP,UAS-GFP/FM7*; *TM3Ser/Sb* (G. Jefferis lab stock), *FRT82B* (RRID:BDSC 2035), *atx2^X1^* (Satterfield et al., 2002), *Me31B::GFP* (RRID:BDSC 51530).

### Immunohistochemistry

#### Adult brains

*Drosophila* adult flies were decapitated, and heads transferred to chilled S2 cell medium (Gibco). The brains were removed from the head capsule and any attached trachea and surrounding tissues were carefully removed. The dissected brains were transferred into a 0.5 ml tube containing 4% paraformaldehyde diluted in PBS containing 0.2% Triton-X100 (PTX) and fixed for 20 min at room temperature on a shaker.

#### Female germline

Mated females (∼5-7 days old) were collected and fed liquid yeast paste for 24h prior to dissection. Ovarioles were isolated from the abdomen of females in PBS, transferred into a 0.5 ml tube containing 4% paraformaldehyde diluted in PBS and fixed for 20 min at room temperature on a shaker.

Fixed samples (adult brains & adult female ovarioles) were washed four times for 15 min each in PTX (PBS + 0.2% Triton), blocked with blocking solution (PTX+ 5% NGS) for 1h and then incubated in primary antibody diluted in blocking solution over night at 4°C. Samples were washed four times for 15 min each in PTX and then incubated in secondary antibodies for 3h at room temperature diluted in blocking solution. Adult brains were mounted in Vectashield Mounting Medium (Vecta labs) on slides using coverslips (thickness No1) as spacers. Female ovarioles in Vectashield were transferred to a microscope slide, dissociated using tungsten needles and sharp forceps and spread out to allow imaging of individual egg chambers.

### Antibodies

Primary antibodies used: mouse rab-Me31B (1:100; Boster), rabbit anti-GFP (1:1000) (Invitrogen), chicken anti-GFP (1:1000) (Abcam). Secondary antibodies used: Alexa 488- and Alexa 555-conjugated anti rabbit and anti-mouse IgG (1:1000) (Invitrogen).

### Image Acquisition

All images were taken on a Zeiss LSM 880 confocal microscope with Airyscan for high resolution imaging. To image Me31B mRNP particles in fixed brain tissue, we used a 63x objective (Zeiss Plan-Apochromate, 1.4 Oil Ph3) and the “digital zoom” software feature of the LSM 880 software (zoom factor between 3 and 4) that allows to scan a region of interest at a higher pixel resolution, constrained of course by the normal ∼300nm resolution limit of light microscopy. Female egg chambers were imaged using a 20x (Zeiss Plan Apochromat 20x/0.8) objective.

### Image analysis

To quantify mRNPs in marked single cells in the adult brain (generated by MARCM) we used a combination of freely available imaging software packages, Fiji (https://fiji.sc) and iLastik (https://www.ilastik.org).

First single channel grey scale stacks for marked mRNP granules were used to train a ‘pixel classification’ in iLastik. After successful training confocal stacks of mRNP granules were segmented and binary images of predicted mRNP granules were exported.

In Fiji a binary mask of single cells marked by GFP was generated by a simple binarization of the GFP channel. This mask was used to subtract the ‘GFP mask’ from the segmented mRNP granules stack leaving only a binary mRNP granules stack that were within the marked cell boundaries.

To quantify mRNPs in single female egg chambers ROIs around the nurse cells were selected with the same size and slice thickness and the image stacks were binarized using the ‘Auto local threshold (Method: Max_Entropy)’ feature in Fiji.

To quantify the number and sizes of mRNP granules in both tissues the Fiji ‘3D Objects counter’ plug in was used. All statistical analysis were performed using GraphPad Prism.

### MARCM to create and visualize single KCs of defined genotype

For the induction of MARCM clones *Drosophila* crosses for both control and experimental were raised at 25°C and the flies were transferred regularly to a new vial for timed egg collections. A heat-shock pulse at 37°C in a water bath for 1-2h was given at optimal time points to induce single cell clones in KCs. For α’/β’KCs 2^nd^ instar larvae (48-60h AEL) and for γKcs early 1^st^ instar larvae (24h AEL) were collected and shocked. Heat-shocked larvae were transferred back to 25°C, raised to adulthood and the adult flies were dissected 1-4 days after eclosion.

### Behavioural assays

Memory experiments were performed at 25°C. Conditioning and testing were carried out in dim red light. *Drosophila* olfactory learning was tested using an olfactory classical conditioning paradigm (Krashes & Waddell, 2011; Tully & Quinn, 1985).

19–24 h before conditioning groups of 60–100 flies were put into vials with 0.75% agar on the bottom for starvation. For appetitive olfactory conditioning, flies were loaded in training tubes lined with water filter papers. After an acclimatization period of 30 s, a first odour (CS-) was presented for 2 min. Then, the odour was removed, and air was allowed to flow for 30 s. Subsequently the animals were transferred to tubes lined with 1% or 2% sucrose filter papers while they were presented with the second odour (CS+) for 2 min. The odour pairs used were ethyl-butyrate [EB] and isopentyl acetate [ISO] or 3-octanol [3-OCT] and 4-methylcyclohexanol [MCH].

For the memory tests, flies were moved to a two-arm choice point where they could choose between the CS+ and CS-for 2 min. After this period, the number of flies within each arm was counted and a preference index was calculated.

PI = (Number of flies in (CS+) – Number of flies in (CS-))/Total number of flies

One memory experiment consisted of two groups of flies with reciprocal conditioning, in which the odour paired with sucrose was exchanged. The preference indices from these two groups were averaged to calculate a performance index (PI).

Short-term memory was tested immediately after conditioning (3 min memory). Flies tested for long-term 24 h memory were put in food vials after conditioning for 6 h and then transferred to starvation vials for 18-20 h until the test.

When conditional expression of the RNAi was required, flies were grown at 18°C and collected after hatching (0–3 d old flies). To induce RNAi expression, flies were moved for 6 d to 29°C. For the starvation period before conditioning and until the test, those animals were kept at 25°C.

All statistical analyses were performed using the GraphPad Prism (https://www.graphpad.com/) and a one-way ANOVA was conducted with a Sidak post-hoc test between the experimental group and the genetic controls.

## RESULTS

### Atx2 is required for the formation of Me31B RNPs in different Kenyon Cell types

Atx2 function has been shown to be essential for the formation of several types of intracellular mRNP granules, including P-bodies and stress granules (Ciosk et al., 2004; Lee et al., 2018; Petrauskas et al., 2024; Singh et al., 2021). Among these, are Me31B/ DDX6-positive mRNP assemblies such as P-bodies and neuronal granules, the latter previously shown to be required for LTH, a form of non-associative long-term memory (Bakthavachalu et al., 2018; Nonhoff et al., 2007; Sudhakaran et al., 2013). Given the potential significance of these assemblies for translational control in neural and non-neural tissue, we performed a MARCM (Mosaic Analysis with a Repressible Cell Marker) (Luo & Wu, 2007) analysis to examine how loss of Atx2 affects Me31B-positive mRNP granules in other cell types. We began by examining mushroom body Kenyon Cells (KCs) known to be involved in well-studied forms of associative memory. MARCM clones of Atx2 were induced following established protocols and methods (Hillebrand et al., 2010; Sudhakaran et al., 2013). Briefly, by using cell-type specific Gal4 lines to drive FLP-recombinase in flies heterozygous for a mutant chromosome, it is possible to induce GFP-marked clones of cells that are homozygously mutated for the protein of interest. This allows to analyse the effect of loss of function of a protein in single cells in an otherwise genetically wild-type background.

We induced Atx2-null clones homozygous for the loss-of-function allele Atx2^X1^ in memory-relevant KCs and examined Me31B granules in these cells. Depletion of Atx2 from α’/β’ and γ KCs caused a clear reduction of Me31B-positive mRNP granules in those neurons [Figure 1]. We compared the total number of mRNP granules per cell [Figure 1 C-I; (C) unpaired t-test p<0.0001, (I) unpaired t-test p<0.0001], average size of RNP granules per cell [Figure 1 D-K; (D) unpaired t-test p<0.0001, (K) unpaired t-test p<0.0001] as well as the number of mRNP granules normalised by the volume of the cell [Figure 1 F-M; (F) unpaired t-test p=0.0018, (M) unpaired t-test p=0.0002]. Each of these analyses revealed significant reduction in both number and size of Me31B mRNP granules in Atx2^X1^ MARCM clones in comparison to control MARCM clones.

**Figure 1.**
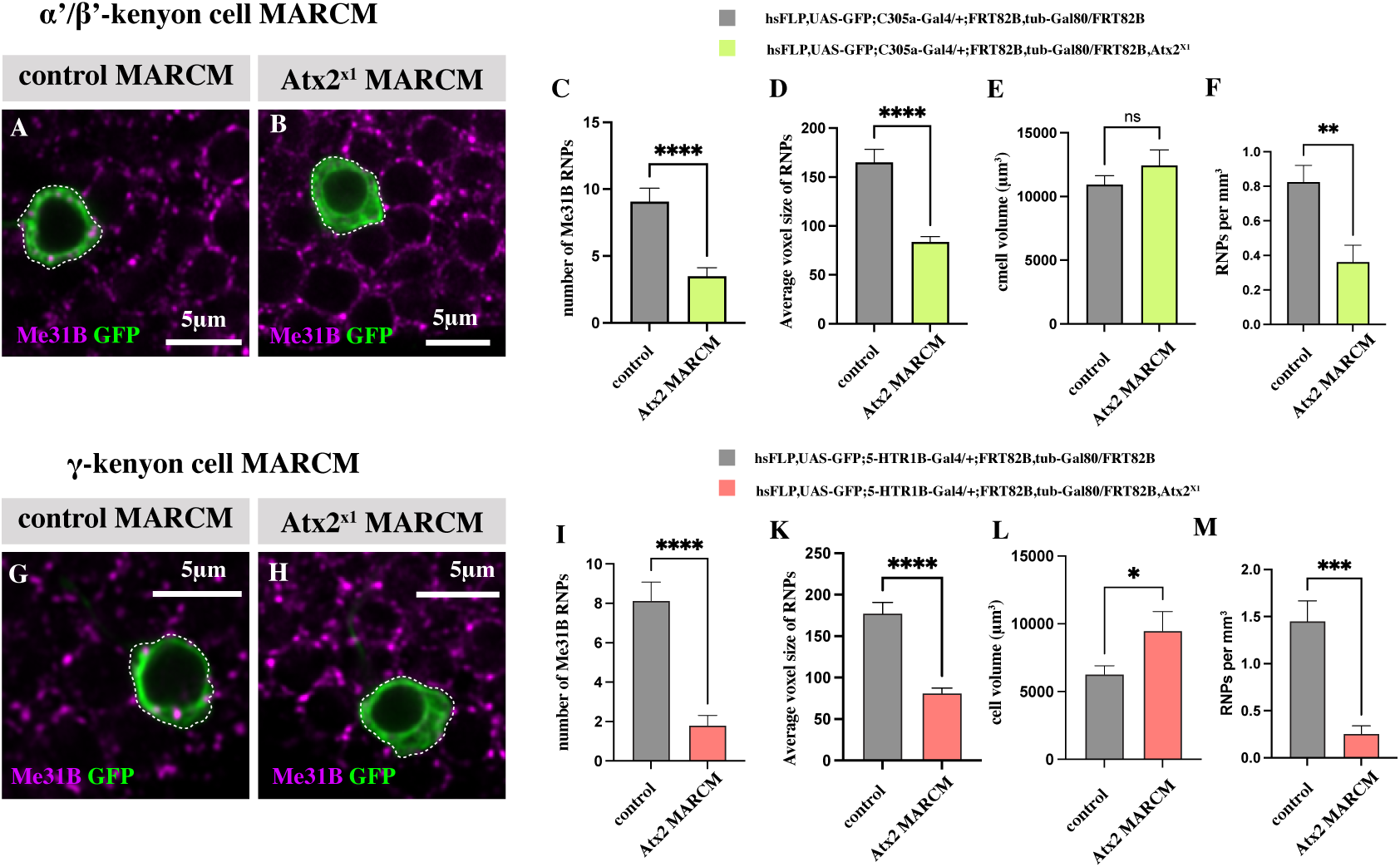
Atx2 is required in α’/β’ and γ KCs for the formation of Me31B-positive mRNP granules. (A) Representative image of control MARCM clones in α’/β’ KCs. (B) Representative image of *atx2^X1^*MARCM clones in α’/β’ KCs. (C) Significant reduction of the total number of Me31B-positive mRNP granules in α’/β’ KCs between control MARCM and *atx2^X1^* MARCM, unpaired t-test p<0.0001. (D) Significant reduction of the average voxel size of Me31B-positive mRNP granules in α’/β’ KCs between control MARCM and *atx2^X1^* MARCM, unpaired t-test p<0.0001. (E) No significant reduction of the volume of α’/β’ KCs cell bodies between control MARCM and *atx2^X1^* MARCM, unpaired t-test p=0.3545. (F) Significant reduction of the number of Me31B-positive mRNP granules normalised to the volume of the cell between control MARCM and *atx2^X1^*MARCM in α’/β’ KCs, unpaired t-test p=0.0018. (C-F) Error bars are ± SEM; ** P<0.01, ****P<0.0001 by unpaired student T-test; control MARCM n=18, *atx2^X1^* MARCM n=20. (G) Representative image of control MARCM clones in γ KCs. (H) Representative image of *atx2^X1^* MARCM clones in γ KCs. (I) Significant reduction of the total number of Me31B-positive mRNP granules in γ KCs between control MARCM and *atx2^X1^* MARCM, unpaired t-test p<0.0001. (K) Significant reduction of the average voxel size of Me31B-positive mRNP granules in γ KCs between control MARCM and *atx2^X1^*MARCM, unpaired t-test p<0.0001. (L) Significant reduction of the volume of γ KCs cell bodies between control MARCM and *atx2^X1^*MARCM, unpaired t-test p=0.0295. (M) Significant reduction of the number of Me31B-positive mRNP granules normalised to the volume of the cell between control MARCM and *atx2^X1^* MARCM in γ KCs, unpaired t-test p=0.0002. (I-M) Error bars are ± SEM; *p<0.05, *** p<0.001, ****p<0.0001 by unpaired student T-test; control MARCM n=14, Atx2^X1^ MARCM n=10.

To further examine whether Atx2 has a broader role in the assembly of Me31B positive mRNP granules across cell types, we tested whether Atx2 in nurse cells of the female germline is required for assembly of well-studied Me31B-positive granules, essential for transport, localization, and timely translation of mRNAs encoding determinants of embryonic pattern (Nakamura et al., 2001; Richter & Lasko, 2011; Voronina et al., 2011). Atx2 knockdown, achieved by expression of either of two different Atx2 RNAi lines during egg development, significantly reduced the efficiency of Me31B-positive mRNP granule assembly in the female germline. This was calculated by analysing the total number of RNP granules [Figure S1 D-F; (D) unpaired t-test p<0.0001, (F) unpaired t-test p=0.0046, (F) unpaired t-test p<0.0001] and the average voxel size of RNP granules [Figure S1 E-G; (E) unpaired t-test p<0.0001, (G) unpaired t-test p=0.0021]. These results, together with previous observations in olfactory projection neurons, local interneurons and cultured S2 cells (Bakthavachalu et al., 2018; Hillebrand et al., 2010; Sudhakaran, Hillebrand et al., 2013), establish Atx2 as a general regulator of Me31B-positive mRNP granules not only across neuronal types, but also across *Drosophila* tissue types.

### Atx2 function is required for the formation of associative LTM

Several translational regulators have been shown to influence a long-term but not a related short-term form of memory (Bolduc et al., 2008; Keleman et al., 2007; McCann et al., 2011; Roselli et al., 2021; Sudhakaran et al., 2013). However, very few such regulators have been tested for roles in more than one type of LTM, let alone in neuronal subtypes required for each form of memory. To examine whether Atx2, previously shown to be required for LTH is also required for long-term appetitive associative olfactory memory, we knocked-down the protein in different Kenyon cell (KC) populations. RNAi expression was restricted to the adult stage by co-expressing a temperature-sensitive inhibitor of Gal4 (Gal80^ts^) under the control of a tubulin promoter (McGuire et al., 2003), along with Gal4-driven RNAi transgenes. In flies carrying *tubGal80^ts^*transgenes, Gal4 is inhibited at low temperatures (18°C) at which Gal80^ts^ is functional, but active when flies are shifted to higher temperatures that are non-permissive for Gal80^ts^ (29°C). After eclosure flies are shifted to 29°C for 6 days, following the animals were moved to a starvation vial at 25°C for 12 hours and then trained for appetitive memory. STM was tested 3 minutes after training while LTM was tested 24 hours after training [Figure 2A].

**Figure 2.**
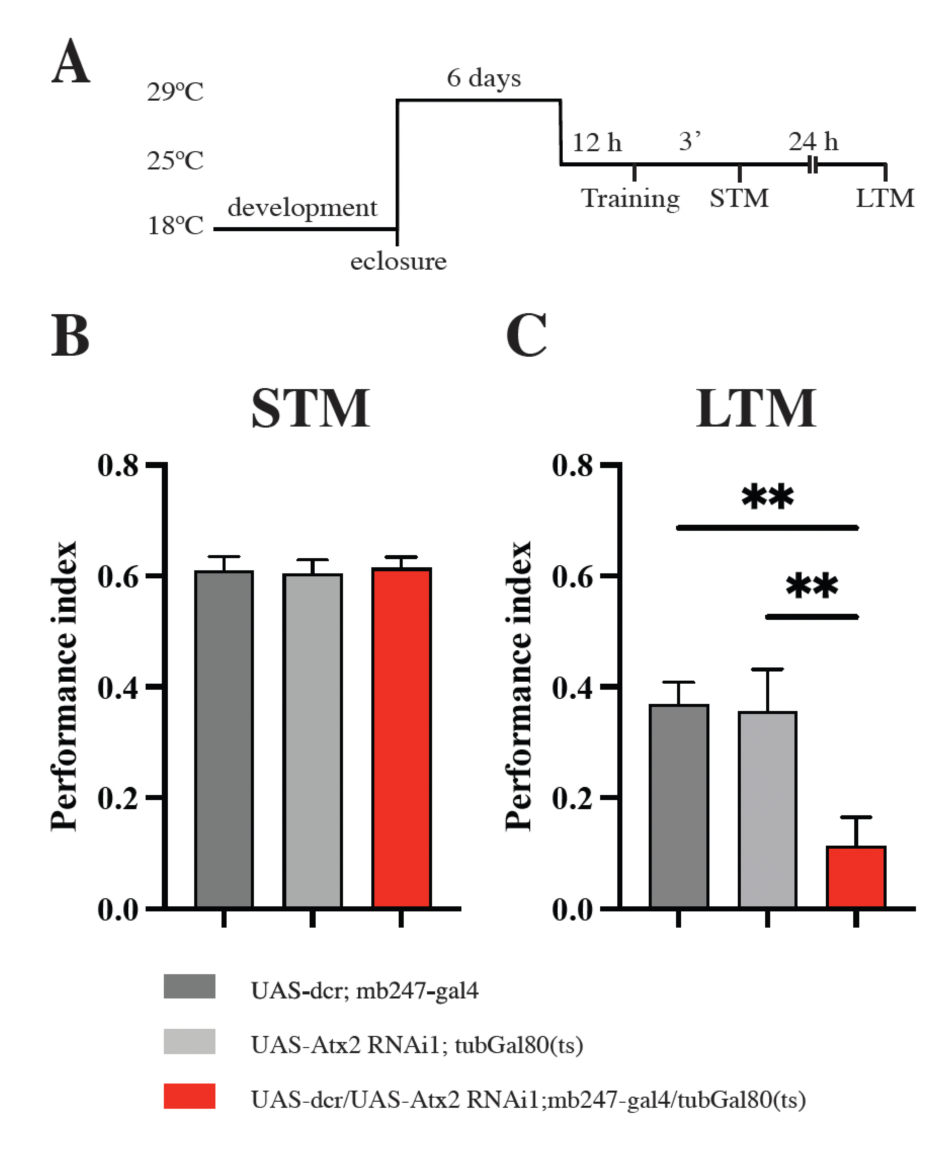
Ataxin-2 (Atx2) protein is required in Kenyon cells (KCs) for expression of olfactory appetitive long-term memories. In adult flies, Atx2 was k/d in a large population of KCs, using the driver *mb247-gal4*. (A) Time course of the experiment. As described in material and methods, the developing animals were kept at 18°C, once eclosed they were collected in group of 60-100 and kept at 29°C to allow RNAi expression for 6 days. Following the animals were moved at 25°C for 12 hours and then trained for appetitive memory. STM was tested 3 minutes after training while LTM was tested 24 hours after training. (B) No difference in olfactory memory performance was found between experimental and genetic control flies at 3 minutes post training (short-term memory), one-way ANOVA p=0.9485; Sidak p=0.9847, p=0.9360. (C) Significant difference in olfactory memory performance was found between experimental and genetic control flies at 24 hours post training (long-term memory), one-way ANOVA p=0.0048; Sidak ** p=0.0069, * p=0.0104. Data are mean performance indices ± SEM; *p<0.05, ** p<0.01 by one-way ANOVA and Turkey’s multiple comparison test; n=12.

Adult-specific expression of Atx2 RNAi using the broad KCs driver, *mb247-gal4* (Aso et al., 2009; Zars et al., 2000) led to a significant decrease in LTM performance in comparison to genetic control flies [Figure 2C; one-way ANOVA p=0.0048], without affecting STM [Figure 2B; one-way ANOVA p=0.9485]. Behavioural control experiments were performed to determine whether the animals could sense and avoid the odorants used for conditioning, as well as for their ability to sense and be attracted to sugar. In all cases, the experimental groups had similar levels of sugar sensitivity, MCH odour and OCT odour avoidance [FigureS2; (A) one-way ANOVA p=0.7436, (B) one-way ANOVA p=0.8015, (C) one-way ANOVA p=0.9889] compared to the controls, indicating a selective role for Atx2 in mediating in KC plasticity processes required for a long-lasting association between behaviourally paired odorants and sugar reward.

To define the specific KCs populations in which Atx2 is required for appetitive LTM we used Gal4 drivers that could target distinct KCs populations: *c739-gal4* for α/β KCs (Akalal et al., 2006), *c305a-gal4* for α’/β’ KCs (Krashes et al., 2007), and *5-HTR1B-gal4* for γ KCs (Aso et al., 2012). Expression of either of two independent Atx2 RNAi lines in α/β [Figure 3A’-B’; (A’) one-way ANOVA p<0.0001, (B’) one-way ANOVA p=0.0227] or α’/β’ [Figure 3 C’-D’; (C’) one-way ANOVA p<0.0001, (D’) one-way ANOVA p=0.0018] KCs, reduced olfactory appetitive LTM measured 24 hours after training without affecting STM [Figure 3 A-B-C-D; (A) one-way ANOVA p=0.6237, (B) one-way ANOVA p=0.0623, (C) one-way ANOVA p=0.9386, (D) one-way ANOVA p=0.0759]. Consistent with prior work showing that γ KCs are mostly involved in short-term appetitive memory rather than in long-term appetitive memory (Cervantes-Sandoval et al., 2013), Atx2 RNAi-expression in γ KCs had no measurable effect on olfactory appetitive LTM [Figure 3 E’-F’; (E’) one-way ANOVA p=0.4982, (F’) one-way ANOVA p=0.5623].

**Figure 3.**
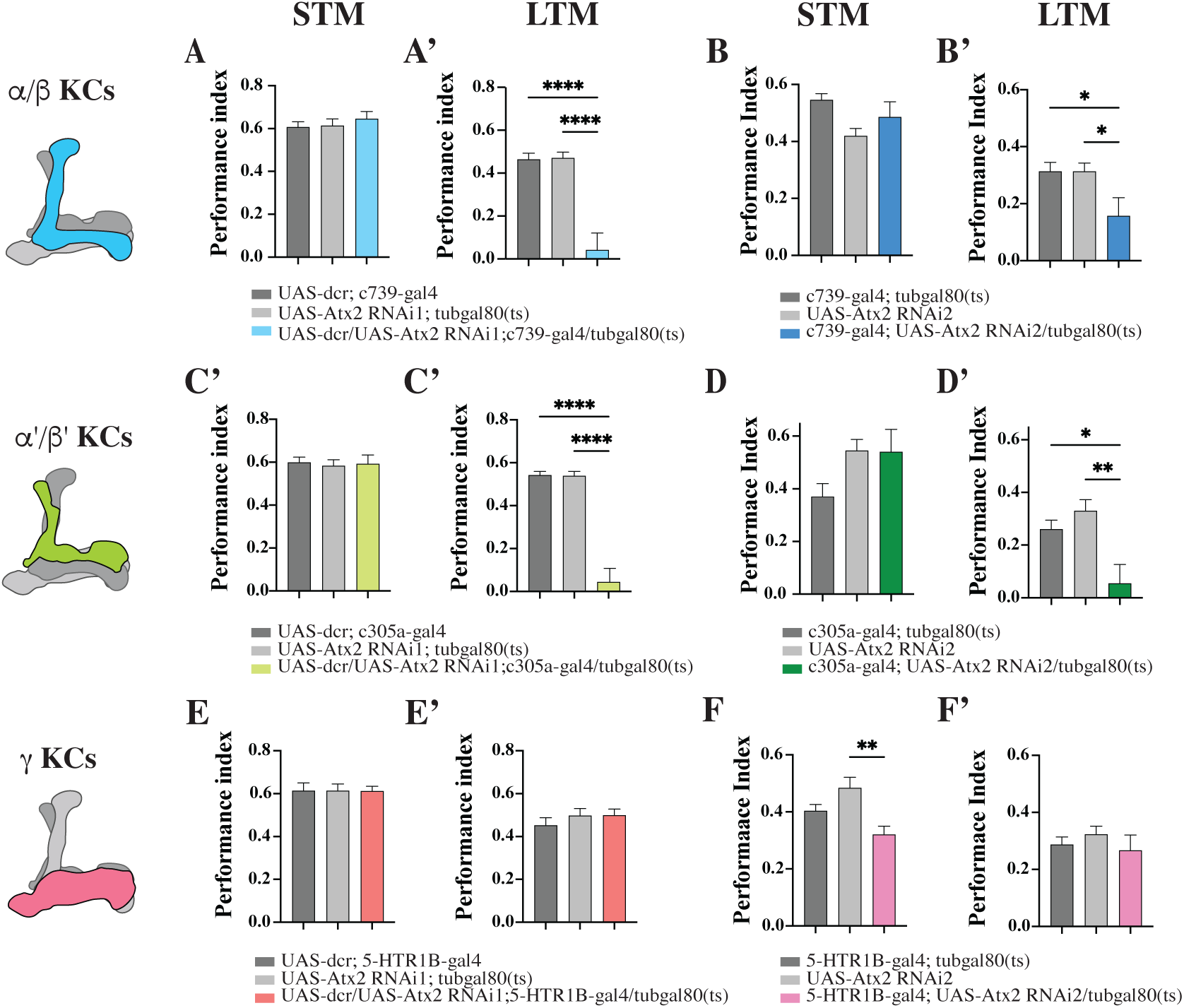
Atx2 is required in α/β and α’/β’ Kenyon cells (KCs) for expression of olfactory appetitive long-term memories. In adult flies, Atx2 was k/d using two RNAis in different KCs populations. (A-A’) Atx2 was k/d using RNAi1 in α/β KCs (*c739-gal4*). (A) No significant difference in STM, one-way ANOVA p=0.6237, Sidak p=0.5983, p=0.7029. (A’) Significant difference in LTM, one-way ANOVA p<0.0001, Sidak **** p<0.0001, **** p<0.0001. (B-B’) Atx2 was k/d using RNAi2 in α/β KCs (*c739-gal4*). (B) No significant difference in STM, one-way ANOVA p=0.0623, Sidak p=0.4382, p=3676. (B’) Significant difference in LTM, one-way ANOVA p=0.0227; Sidak * p=0.0311, * p=0.0311. (C-C’) Atx2 was k/d using RNAi1 in α’/β’ KCs (*c305a-gal4*). (C) No significant difference in STM, one-way ANOVA p=0.9386, Sidak p=9857, p=9744. (C’) Significant difference in LTM, one-way ANOVA p<0.0001, Sidak **** p<0.0001, **** p<0.0001. (D-D’) Atx2 was k/d using RNAi2 in α’/β’ KCs (*c305a-gal4*). (D) No significant difference in STM, one-way ANOVA p=0.0759, Sidak p=0.1077, p=0.9983. (D’) Significant difference in LTM, one-way ANOVA p=0.0018; Sidak * p=0.0149, ** p=0.0012. (E-E’) Atx2 was k/d using RNAi1 in γ KCs (*5-HTR1B-gal4*). (E) No significant difference in STM, one-way ANOVA p=0.9991, Sidak p=0.9991, p=0.9992. (E’) No significant difference in LTM, one-way ANOVA p=0.4982, Sidak p=0.5112, p=0.9992. (F-F’) Atx2 was k/d using RNAi2 in γ KCs (*5-HTR1B-gal4*). (F) Significant difference in STM between one control and the experimental line, one-way ANOVA p=0.031, Sidak p=0.1675, p=0.0015. (F’) No significant difference in LTM, one-way ANOVA p=0.5623, Sidak p=9171, p=5023. Data are mean performance indices ± SEM; * p<0.05, ** p<0.01, ****p<0.0001 by one-way ANOVA and Turkey’s multiple comparison test; n=8-12.

Thus, Atx2 function is required not only in olfactory PNs and LNs for long-term habituation (Bakthavachalu et al., 2018; McCann et al., 2011; Sudhakaran et al., 2013), but also in KCs subtypes for long-term olfactory associative memory.

## DISCUSSION

This study has investigated the role of the translational regulator Atx2 in mRNP granules assembly and in long-term memory formation. We can draw two main conclusions from these experiments: (a) Atx2 contributes to the assembly of different types of Me31B-positive granules in multiple neuronal and non-neuronal cells; and (b) Atx2 is a fundamental component of plasticity processes that is required for long-term forms of memory.

### Atx2 regulates the formation of Me31B-positive mRNP granules in different tissues

Ribonucleoprotein (mRNP) granules are a diverse group of RNA–protein assemblies. Distinct types of mRNP granules can have different localizations, RNA and protein compositions, as well as different roles in RNA metabolism, biological function and disease (Protter & Parker, 2016; Tauber et al., 2020).For instance, nucleolar granules are sites of rRNA biogenesis, stress granules mediate aspects of translational arrest and metabolic change in response to oxidative, radiational, nutritional or viral stress (Buchan & Parker, 2009; Tauber et al., 2020) and cytoplasmic P-bodies contain RNA decay machinery that mediate mRNA turnover (Parker & Sheth, 2007; Stoecklin & Kedersha, 2013). In addition, there are tissue-specific and highly specialized granules including germline and neuronal granules thought to operate as hubs for posttranscriptional regulation of gene expression required for germ cell and early embryonic development, neuronal asymmetry, and localized mRNA translation (Kato & Nakamura, 2012; Trcek et al., 2015; Voronina et al., 2011). While some types of neuronal granules are used for transport of translational arrested mRNAs, other related types may be required for local storage and translational control of synaptic mRNAs – a process that is required for long-term forms of synaptic plasticity (Bakthavachalu et al., 2018; Grzejda et al., 2022; Krichevsky & Kosik, 2001). What remains unclear is the range of neuronal granule types that mediate local translational control, as well the diversity of mechanisms and their dynamics *in vivo*.

We focus here on a class of neuronal RNP granule that contains Me31B, a DEAD-Box RNA helicase homologous to human DDX6, and requires Atx2 for their normal assembly. While such granules exist in multiple neuronal subtypes, it is important to note that because different granule types can share common components and regulators (Gruidl et al., 1996; Tindell et al., 2020), there could be more than one subtype of Me31B granule targeted in our experiments. For instance, while germline and neuronal Me31B-positive granules regulate the translation of respective target mRNAs and depend on Atx2 for their assembly, they are distinctive in composition and biological function [Figure 1, Figure S1]. Nonetheless our experiments unequivocally demonstrate that Atx2 is necessary for the assembly of Me31B foci in multiple neuronal types, not only in PNs and LNs of the antennal lobe implicated in long-term habituation., but also crucially in KCs known to encode long-term associative memory.

### Atx2 dependent Translational Control in Long-Term memory

Our data support models for two aspects of molecular mechanisms that underlie long term memory formation in the mushroom body circuit. First, they provide new support for a model in which LTM of odour-sugar association requires long-term plasticity in synapses two different types of KCs. Second, by demonstrating that Atx2 is required for long-term plasticity in different brain regions, that Atx2 dependent translational control is broadly required for different forms of long-term memory.

The circuit underlying LTM allows experience-induced synaptic plasticity to be studied in the context of behavioural memory. Previous and extensive work on *Drosophila melanogaster* has pointed to the KCs as the main coding centre for associative olfactory memories in the fruit fly (Dubnau & Tully, 2001; Heisenberg et al., 1985; McGuire et al., 2001; Trannoy et al., 2011). The observation that synaptic outputs from both α/β and α’/β’ KCs populations are required for LTM is suggestive of plasticity being required in multiple circuit locations for LTM (Cervantes-Sandoval et al., 2013; Owald et al., 2015; Trannoy et al., 2011). Consistent with this, new gene expression in the MB (Jones et al., 2018; Kramer et al., 2011; Petruccelli et al., 2020; Widmer, Bilican, et al., 2018) and *de novo* protein synthesis in KCs has been shown to be required for LTM of associative conditioning (Chen et al., 2012; Hirano et al., 2013; Yu et al., 2006). Function of multiple translation control molecules such as dFMR/ FMRP was shown to be required in KCs for the formation of aversive LTM (Bolduc et al., 2008).

More detailed studies of the spatial requirement for gene expression showed that activity of the transcriptional regulator CREB2 is needed in two specific subpopulations of the KCs, α/β and α’/β’ for consolidating LTM of appetitive conditioning (Krashes & Waddell, 2008; Widmer, Fritsch, et al., 2018; Hirano et al., 2016; Yin et al., 1994). The hypothesis that this indicates a need for protein-synthesis dependent plasticity in these two classes of KCs for LTM formation, is now further supported by our observation that olfactory appetitive LTM similarly requires the translational and mRNP granule regulator Atx2 in α/β and in α’/β’ adult stage KCs [Figure 3 A’-D’] without affecting STM or olfactory responses [Figure 3 A-D].

While our data do not formally demonstrate that Atx2 functions in local translation of synaptic mRNAs in KCs, it demonstrates though a specific requirement for Atx2 for cell type-specific LTM but not STM, which provides a strong argument for local translation of mRNAs being the underlying mechanism by which these proteins regulate long-term plasticity of α/β and in α’/β’ KCs *in vivo*.

Future studies will be needed to show the translation *in vivo* of Atx2 target mRNAs and will aim to investigate whether Atx2 could be required in other neuronal populations, which have been implicated in translational regulation of LTM such as mushroom body output neurons or DAL (Chen et al., 2012; Crocker et al., 2016; Hirano et al., 2013; Wu et al., 2017) for LTM formation. However, taken together with prior work showing that long-term olfactory habituation requires Atx2 in specific subgroups of projection neurons and local interneurons (Bakthavachalu et al., 2018; McCann et al., 2011; Sudhakaran, et al., 2013), these findings support a central and broad role for Atx2 in the control of protein synthesis dependent long-term plasticity across brain systems in *Drosophila* and potentially across species.

## ACKNOWLEDGMENTS

We thank the members of the Ramaswami lab for useful discussions. CR was funded by the Irish Research Council Postgraduate scholarship. MR acknowledges support from a Wellcome Trust-SFI-HRB Investigator grant, an Irish Research Council Laureate Award and an SFI Frontiers for the Future grant.

---

**Figure S1.**
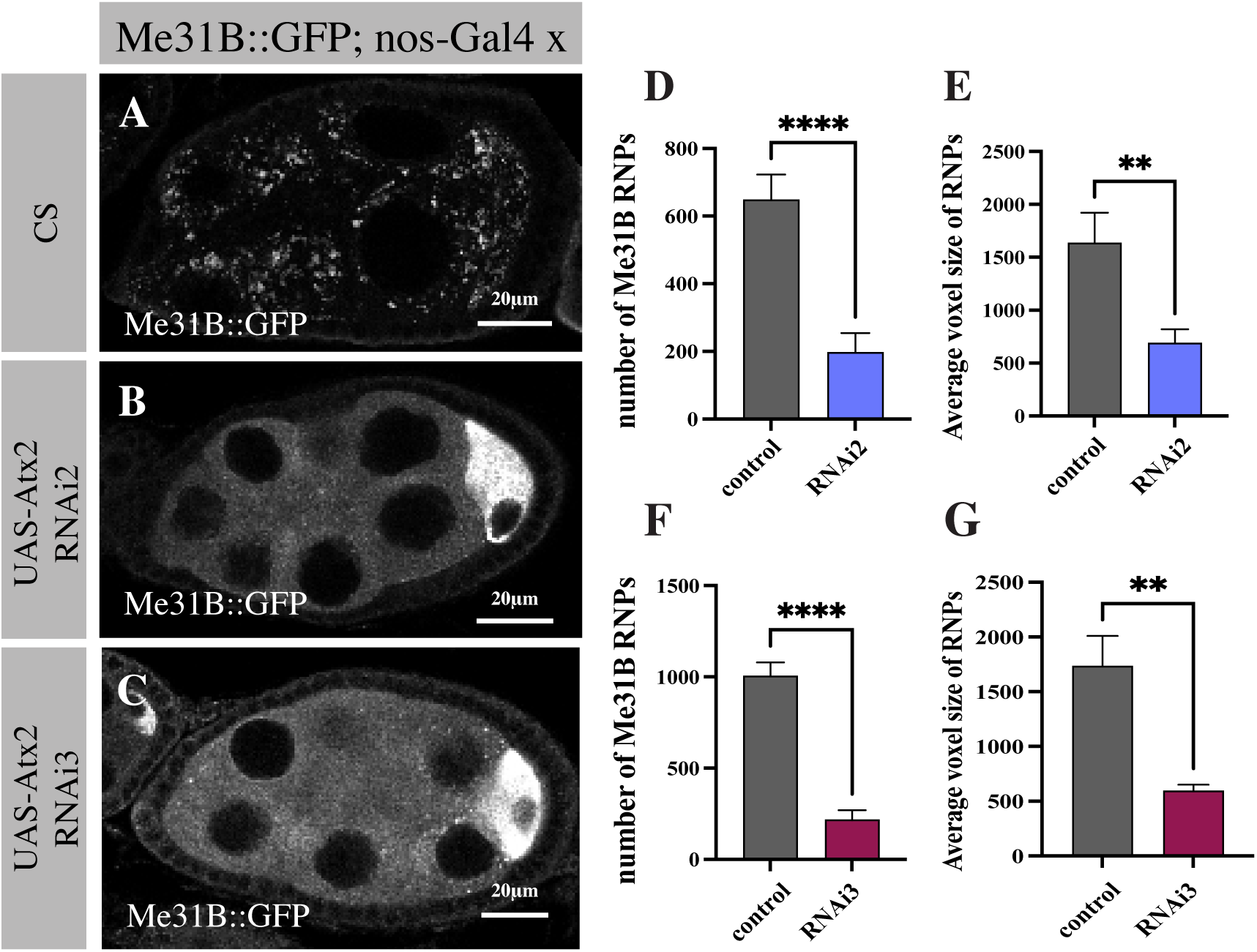
Atx2 protein is required in female egg chamber nurse cells for the formation of Me31B-positive mRNP granules. (A) Representative image of Me31B expression (marked by a Me31B::GFP protein trap line (Buszczak et al., 2007) in a stage 6 egg chamber). (B) Depletion of Atx2 by expression of UAS RNAi2 and (C) UAS RNAi3, expressed by nos-Gal4 abolishes Me13B-positive mRNP granules in nurse cells. (D) Quantification of Me31B-positive mRNP granules numbers in control and Atx2 k/d using RNAi 2, unpaired t-test p<0.0001. (E) Quantification of Me31B-positive mRNP granules average voxel size in control and Atx2 k/d using RNAi 2, unpaired t-test p=0.0046. (F) Quantification of Me31B-positive mRNP granules numbers in control and Atx2 k/d using RNAi 3, unpaired t-test p<0.0001. (G) Quantification of Me31B-positive mRNP granules average voxel size in control and Atx2 k/d using RNAi 3, unpaired t-test p=0.0021. Both Atx2 RNAi lines show a significant reduction of Me31B RNP numbers and size quantified in nurse cells of eggchambers at the same developmental stages. Error bars are ± SEM; *** p<0.01, ****p<0.0001 by unpaired student T-test; control eggchambers n=10, Atx2-RNAi2 n=12, Atx2-RNAi3 n=9.

**Figure S2.**
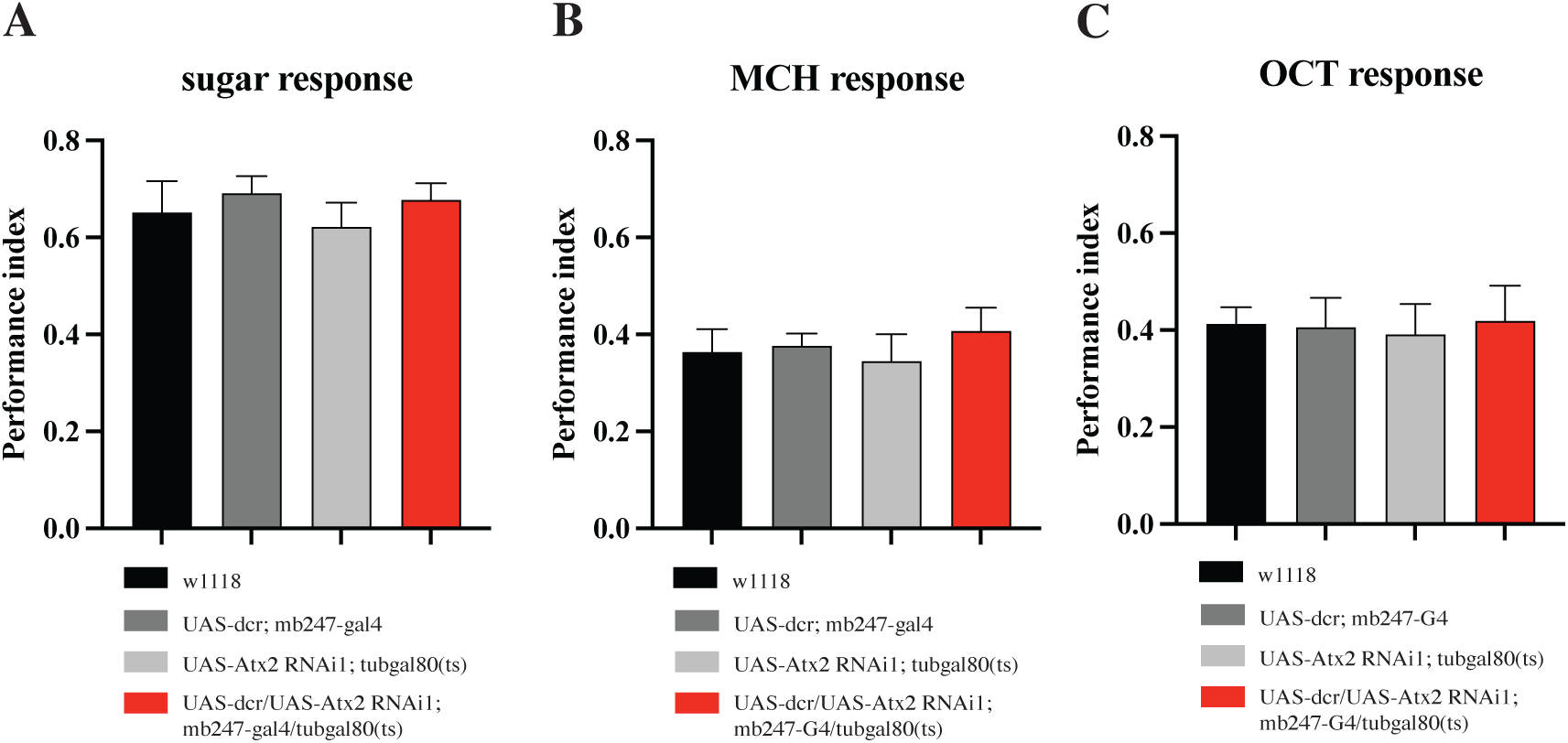
Atx2 RNAi expression in Kenyon cells does not affect sugar attraction or odour avoidance. In adult flies, Atx2 was k/d in a large population of KCs, using the driver mb247-gal4. (A) The sugar attraction test was performed with a sugar stripe compared with a water stripe in the T-maze. No significant difference was found between experimental and genetic control flies, one-way ANOVA p=0.7436, Sidak p=0.9732, p=9966, p=7907. (B) Olfactory avoidance was measured to test the ability of the experimental and genetic control flies to sense and avoid the MCH odour used for conditioning. No significant difference was found between experimental and genetic control flies, one-way ANOVA p=0.8015, Sidak p=0.8797, p=0.9512, p=7074. (C) Olfactory avoidance was measured to test the ability of the experimental and genetic control flies to sense and avoid the OCT odour used for conditioning. No significant difference was found between experimental and genetic control flies, one-way ANOVA p=0.9889, Sidak p=0.9998, p=9980, p=9826. Error bars are ± SEM.

